# The Virtual Mouse Brain: a computational neuroinformatics platform to study whole mouse brain dynamics

**DOI:** 10.1101/123406

**Authors:** Francesca Melozzi, Marmaduke M. Woodman, Viktor K. Jirsa, Christophe Bernard

## Abstract

Connectome-based modeling of large-scale brain network dynamics enables causal in silico interrogation of the brain’s structure-function relationship, necessitating the close integration of diverse neuroinformatics fields.

Here we extend the open-source simulation software The Virtual Brain to whole mouse brain network modeling based on individual diffusion Magnetic Resonance Imaging (dMRI)-based or tracer-based detailed mouse connectomes. We provide practical examples on how to use The Virtual Mouse Brain to simulate brain activity, such as seizure propagation and the switching behavior of the resting state dynamics in health and disease.

The Virtual Mouse Brain enables theoretically driven experimental planning and ways to test predictions in the numerous strains of mice available to study brain function in normal and pathological conditions.

## Introduction

Dedicated software environments are available to simulate detailed neuronal dynamics such as Neuron, Genesis and MOOSE, which model the complex dendrite geometry, reaction-diffusion processes and receptor distributions of individual neurons and smaller networks (Hines and Carnevale, 1997). To simulate larger networks, neuron models are reduced to point neurons (Goodman and Brette, 2009, Brette and Gerstner, 2005, Izhikevich et al., 2003). However, scaling up for detailed models beyond an entire cortical column (Markram, 2012) and a few brain regions becomes quickly intractable even for networks of point neurons. Even though neuromorphic computation offers interesting alternatives for the future (SpiNNaker, Project, BrainScales, project), macroscopic modeling using neural population approaches is the only viable whole brain network modeling strategy nowadays.

The Virtual Brain (TVB) is an open-source simulation software designed to model whole-brain network dynamics, where the network’s connectivity is based on diffusion MRI-based individual connectomes or adaptations of more precise primate connectomes (Leon et al., 2013). TVB comprises several generative neural population models, defined in physical 3D space and constrained by anatomy, allowing simulating neuroimaging signals (such as magneto-and electroencephalography (MEG, EEG), or functional MRI (fMRI)). Whole brain dynamics can also be manipulated in TVB, e.g. via stimulation. TVB provides a large set of tools for visualization and data analysis (Sanz-Leon et al., 2015). As such, TVB provides a conceptual framework to interpret neuroimaging data, offering promising diagnostic and therapeutic perspectives (Jirsa et al., 2016, Proix et al., 2017). Other groups demonstrate converging results using similar large-scale brain modeling approaches (Hutchison et al., 2013, Sinha et al., 2016). However, very few model predictions can be experimentally tested in humans for obvious ethical reasons. Thus, assessing causality and extracting general principles of brain dynamics in health and disease remain a challenge.

Rodent research enabled major advances in our understanding of brain function and dysfunction, but mostly at the microscopic scale. The advent of new generations of MRI machines now gives access to detailed anatomical, structural and functional information at the whole rodent brain scale (Stafford et al., 2014), thus providing a formidable opportunity to explore general principles of whole brain dynamics. Indeed, hypotheses can be tested and causality can be assessed in the numerous transgenic mouse lines that have been generated to study neurological disorders and to manipulate neuronal networks (e.g. with optogenetics and pharmacomogenetics). However, a conceptual framework is needed to interpret neuroimaging data and generate testable hypotheses. Such framework would considerably accelerate our understanding of the mechanisms controlling and affecting whole brain dynamics.

Here we present The Virtual Mouse Brain (TVMB), the first connectome-based simulation platform to study large-scale mouse brain dynamics.

TVMB is an extended version of TVB, adapted to the mouse brain to enable its virtualization. It inherits from TVB all the already validated simulators to generate brain network activity, as well as analysis and visualization tools.

In what follows, we will show how the platform can be used to virtualize not only individual mouse brains (based on diffusion MRI connectome) but also to construct very detailed connectome-based models using tracer data generated by the Allen Institute for Brain Science (Oh et al., 2014).

As a worked example to show how the platform can be used to generate predictions or interpret data, we will simulate resting state dynamics in a control and “epileptic” mouse, and seizure propagation. We will also show how to integrate TVMB in a research project in which theoretical and experimental approaches benefit from one another.

## Materials and methods

All the methods discussed in what follows are implemented in TVB and freely available to the community. In supplementary materials are contained all the instructions, materials and codes necessary to reproduce the same results presented in the paper; in particular Tutotial TVMB contains all the general information to run TVMB.

### The Allen Connectivity Builder

The Allen Connectivity Builder is a pipeline that we have designed in order to build a complete mouse connectome based on tracer information.

Specifically we define the link between two brain regions according to the anterograde tracing information provided by the Allen Institute of Brain Science and presented in the work of Oh et al. (2014). In the latter, the axonal projections from a given region are mapped by injecting in adult male C57Bl/6J mice the recombinant adeno-associated virus, which expresses the EGFP anterograde tracer. The tracer migration signal is detected with a serial two-photon tomography system. This approach is repeated systematically in order to collect the information on the tracer migration from several injection sites in the right hemisphere to target regions in both ipsilateral and contralateral hemispheres; for each injection sites several experiments are run and distinct measures are accomplished. The Allen Institute provides its data through an internet-accessible interface, namely the Allen Software Development Kit (Allen SDK), from which TVB, through the Allen Connectivity Builder interface, is able to obtain a volumetric atlas as well as the raw experimental information necessary to build complete mouse brain connectomes. The platform allows to chose the main characteristics of the connectome; specifically the user can set:

1. the resolution of the grid volume in which the data are registered (25 *μm*, 50 *μm*, 100 *μm*).
2. The definition of the connection strength between source region *i* and target region *j*. Specifically the connection strength can be defined as:

–The detected projection density (the number of detected pixels in the target region normalized on the total number of pixels belonging to that region).
–The detected projection energy (the intensity of detected pixels in the target region normalized on the total number of pixels belonging to that region).
–The ratio between the projection density, defined as explained above, and the injection density of the source region (the number of infected pixels in the source region normalized on the total number of pixels belonging to that region). It is possible to choose the characteristics of the brain areas to be included in the parcellation using the two following criteria:
3. Brain areas where at least one injection has infected more than a given threshold of voxels. This kind of selection ensures that only the data with a certain level of experimental relevance are included in the connectome (Oh et al., 2014).
4. Only brain areas that have a volume greater than a given threshold can be included.

The pipeline, once downloaded the raw data from the Allen dataset, cleans the data in order to obtain a set of experiments in which the injection structures are exactly the same as the target structures and vice versa; this step ensures that the connectome will be a square matrix. Then, the pipeline excludes from the experimental set the area that do not fulfill the criteria set by the user (minimum volume (3) and minimum number of voxels infected (4)).

The experiments of the Allen Institute consider source regions always located in the right hemisphere, thus we build a complete structural connectivity matrix, taking the mirror image of the right hemisphere to build the left one. Therefore, if we divide the SC matrix in four blocks R-R, R-L, L-R and L-L (clockwise order starting from upper left), we will have the symmetries R-R = L-L and R-L = L-R. This assumption is justified by the fact that the mouse brain shows a high degree of lateral symmetry (Calabrese et al., 2015).

The connection strength between a given region and another one is averaged across all the experiments that use as source and target regions those particular brain areas. The Allen Connectivity Builder approximates the length of the tracts as the Euclidean distance between the centers of the brain regions; the latter are calculated using the volume built from the Allen SDK.

Finally the Allen Connectivity Builder create a region volume mapping, i.e. a 3D matrix which represents the volume of the mouse brain, by modifying the annotation volume downloaded from the Allen SDK. In particular the volume is built so that the entries of the 3D volume matrix range from -1 (background) to N-1, where N is the total number of areas in the connectivity: entries in the volume equal to *i* − 1 label the brain region whose incoming and outgoing connections are organized in the *i*-th row and *i*-th column of the connectivity matrix.

The volumes and the connectivities used in the present work have a resolution of 100 *μm* and each connection strength is defined as the ratio between projection density and injection density. The areas included in the parcellation have a volume greater than 2 *mm*^3^ and they have more than 50 voxels infected in at least one injection experiment. The connectome obtained is in Connectivity.zip folder (Supplementary materials); the instruction to obtain it through the TVB GUI and the Jupiter interface are respectively in TVMB_tutotial.pdf and Create_connectome_Allen.ipynb.

### Resting state dynamics

#### Brain model

The mean activity of each brain region, composing the mouse brain network, is described by the reduced Wong Wang model (Wong and Wang, 2006). In this approach the dynamics of a brain region is given by the whole dynamics of excitatory and inhibitory populations of leaky integrate-and-fire neurons interconnected via NMDA synapses. In this work we take into account this model with a further reduction performed in Deco et al. (2013): the dynamics of the output synaptic NMDA gating variable *S* of a local brain area *i* is strictly bound to the collective firing rate *H_i_*. The resulting model is given by the following coupled equations:

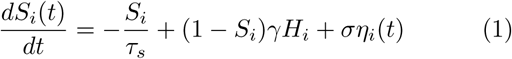

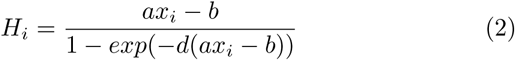

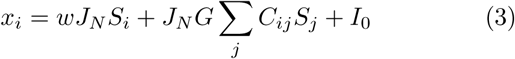
where *x_i_* is the synaptic input to the *i*-th region. *γ* is a kinetic parameter fixed to 0.641, *τ*_s_ is the NMDA decay time constant and its value is 100 ms; *a, b* and *d* are the parameters of the input and output function *H* and are respectively equal to 270 *nC^−^*^1^, 108 Hz, 0.154 s. *J_N_* = 0.2609 nA is an intensity scale for the synaptic input current. *w* is the local excitatory recurrence and *I*_0_ is the external input current. The value of the local excitatory recurrence, *w*, and the external input current, *I*_0_, are set respectively to 0.3 nA and 1 in order to enrich the non-linearity of the dynamics of each brain region; this has an impact on the global network by introducing attractors that are not in trivial relation with the anatomical connectivity, as predicted in the study of Hansen et al. (2015); we refer to this model as the *enhanced non-linearity Mean-Field Model* (eMFM). The implementation of the eMFM in a brain network offers the chance to study the non-stationary features of the functional connectivity patterns.

*G* is the coupling strength i.e. a scalar parameter which scales all the connection strengths *C_ij_* without altering the global topology of the network. The value of G together with the value of the noise amplitude *σ* of the normally distributed stochastic variable *η_i_*, are tuned resoectively to 0.096 and 5.1 · 10^−3^ in order to initialized in the optimal regime to simulate the resting state activity, i.e. the regime where the system explores different states (Deco et al., 2011,Deco and Jirsa, 2012, Hansen et al., 2015).

#### Integration scheme and BOLD signals

Model equations are numerically solved using the Euler integration method with a fixed integration step of 0.1 ms.

Simulated BOLD signal is obtained by converting the simulated neural activity using the Balloon-Windkessel method (Friston et al., 2000) using the default value implemented in The Virtual Brain (Sanz-Leon et al., 2015). The BOLD time-series are down-sampled to 2 s and 20min total length.

#### Functional connections

Functional connections in the simulated time-series are explored from both spatial and temporal point of views using, respectively, the functional connectivity (FC) and the functional connectivity dynamics (FCD).

The *ij*-th element of the FC matrix is calculated as the Pearson correlation between the BOLD signal of the brain region *i* and of the brain region *j*.

To estimate the FCD, the entire BOLD time-series is divided in time windows of a fixed length (3 min) and with an overlap of 176 s; the data points within each window centered at the time *t_i_* were used to calculate FC(*t_i_*). The *ij*-th element of the FCD matrix is calculated as the Pearson correlation between the upper triangular part of the *FC* (*t_i_*) matrix arranged as a vector and the upper triangular part of the FC(*t_j_*) matrix arranged as a vector. In order to observe signal correlations at frequency greater than the typical one of the BOLD signals, the sliding window length is fixed to 3 min, since, as demonstrated by Leonardi and Van De Ville (2015), the non-spurious correlations in the FCD are limited by high-pass filtering of the signals with a cut-off equal to the inverse of the window length.

The FCD matrix allows identifying the epochs of stable FC configurations as blocks of elevated inter-*FC*(*t*) correlation; these blocks are organized around the diagonal of the FCD matrix (Hansen et al., 2015).

#### FCD segmentation: Spectral Embedding

In order to identify the epochs of stable FC configurations, we used the spectral embedding method, that permits to group together the nodes of the FCD, i.e. the different time windows, in clusters.

The spectral embedding is a general cluster technique founded on the possibility to map the nodes of the network in the Euclidean space such that the Euclidean distance between the nodes in the space corresponds with the distance between the nodes in the network.

In order to implement this idea, it is necessary to define the notion of *distance* between nodes in a network; this is made introducing the concept of the commute distance *c_ij_* between the nodes *i* and *j* that is defined as the expected number of steps in a random walk starting to travel from node *i* to node *j*, and back (Von Luxburg, 2007).

To mathematically define *c_ij_* is necessary to introduce some quantity. Let us consider a graph that has an adjacency matrix *W*, i.e. a matrix whose element *w_ij_* is the weight of the link between node *i* and *j*, that in our case is the FCD matrix; it is possible to define the laplacian of the graph as:

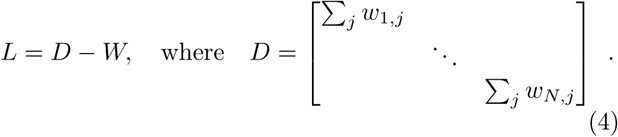

Let us denote with |*e_i_*〉 the eigenvector *i* of the Laplacian; if the matrix *U* is the matrix whose columns are the eigenvectors of *L*, and Λ the diagonal matrix with the eigenvalues *λ_i_* on the diagonal, thus it is possible to decompose the Laplacian as: *L* = *U*Λ*U^T^*.

The generalized inverse of the Laplacian is defined as *L*^†^ = *U*Λ^†^*U^T^*, where Λ^†^ is the diagonal matrix with on the diagonal 1/*λ_i_* when *λ_i_* is different from zero, otherwise zero. Thanks to *L*^†^ it is possible to express the commute distance between node *i* and *j* as:

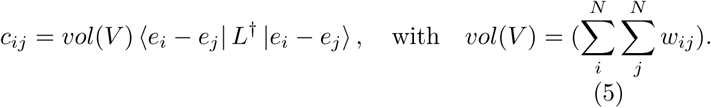

The variable |*z_i_*〉 maps the vertex υ*_i_* in the Euclidean space (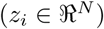) such that the Euclidean distance between node *i* and *j* is equal to the commute distance *c_ij_* of the nodes in the graph if and only if:

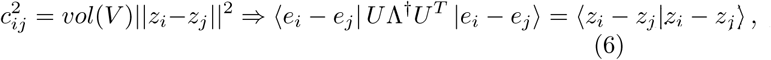
from which it follows that 〈*z_i_*| corresponds to the *i*–th row of the matrix 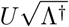.

In Figure 1-1 (Supplementary materials) the nodes of the FCD are plotted in the Euclidean space 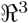 mapping only the first two components of 〈*z_i_*|.

**Figure 1:**
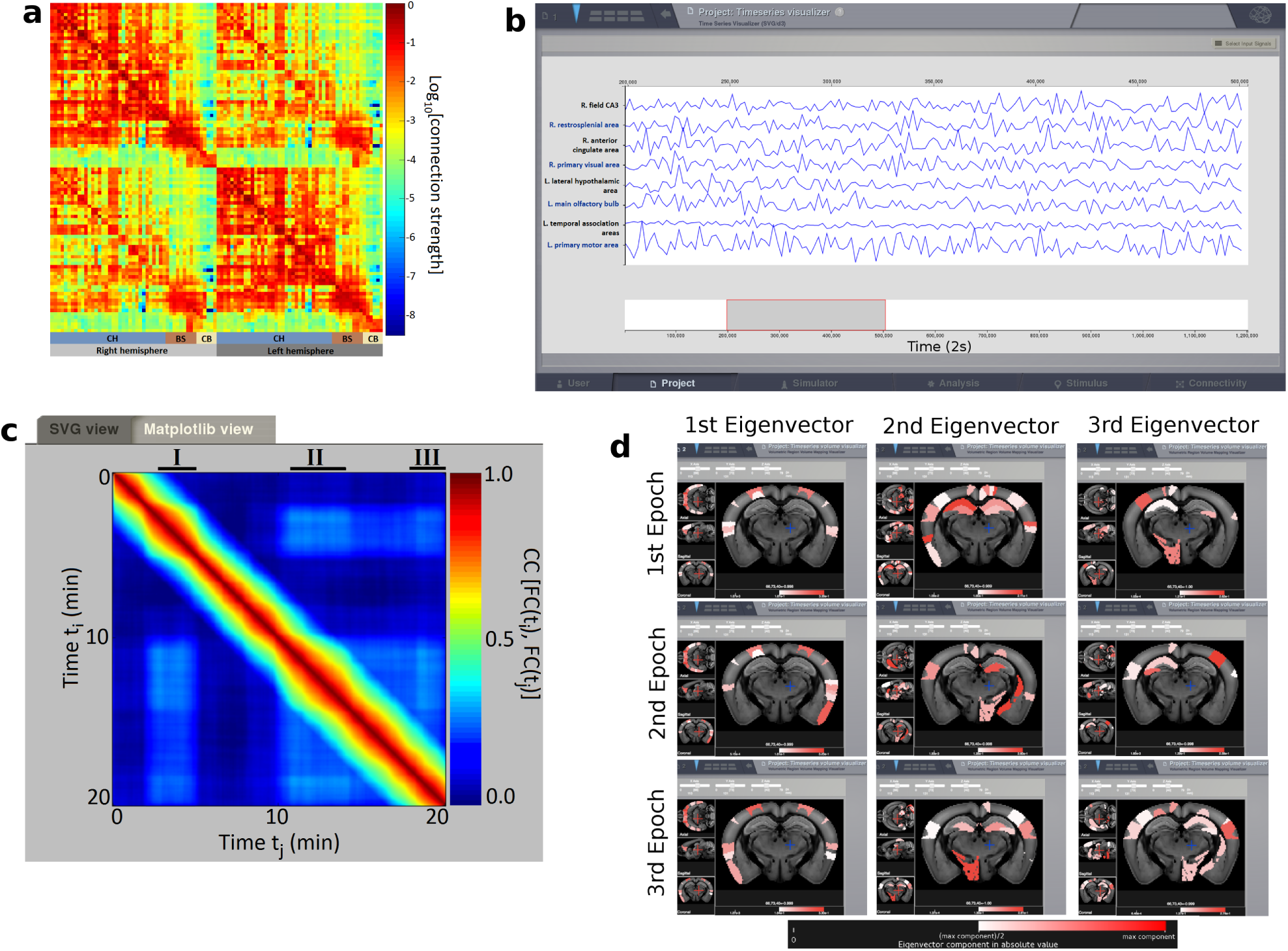
(a) We used the Allen Connectivity Builder to build the structural connectivity matrix. The color map represents the connections strengths with a base-ten logarithmic scale. The resolution of the grid is 100 *μm* the weights of the matrices are defined as the ratio between the projection and the injection density; all the areas in the parcellation have at least one injection experiment that has infected more than 50 voxels in those areas; the matrix contains only regions with a volume greater than 2 mm^3^. (b) Simulated resting state BOLD time-series using the connectome built in (a), and the eMFM to model the dynamics of each brain area. (c) FCD matrix obtained from the time-series. The three black segments (I, II and III) correspond to epochs of stability of the FCD identified with the spectral embedding technique. (d) Functional hubs detected in silico mapped on brain sections using the brain region volume visualizer. Images in the same row represent the plotting of the eigenvectors components, in absolute value, of the FC belonging to the same epoch. Images organized in different columns refer to eigenvectors belonging to different eigenvalues of the matrices. The scale used allows highlighting only the brain area associated to component of the eigenvector greater than the half of the maximum component. Such scale permits to efficiently visualize the relative difference between eigenvectors. According to our definition (see method section), the areas with warm colors are the hub regions of the brain network defined by the FC matrices calculated over the relative epoch; the importance of each hub region is proportional to the corresponding eigenvalue. The instructions and the codes to obtain the results in figure are respectively in Tutorial 1-1 and code 1-1, code 1-2 (Supplementary materials).

#### Functional hubs

The functional connectivity matrix of each epoch defines a functional network; for each functional network, we identify the hub regions with an approach analogous to the one used in graph theory for defining the eigenvector centrality of a network node (Newman, 2008).

Let us define the functional centrality *φ*^(^*^i^*^)^ of a brain region *i* as the sum of the functional centralities of the neighboring brain regions weighted on the functional connection strength *fc_ij_*:

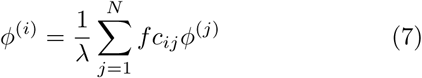
where λ is a constant. Defining the vector 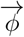 as the column vector whose components are the functional centrality of each network region, we can rewrite the previous equation in matrix form

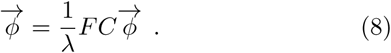

It is simple to notice that 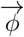 is the eigenvector of the functional connectivity matrix associated with the eigenvalue λ. Since the FC is a real symmetric matrix (thus diagonalizable), we can decompose it as:

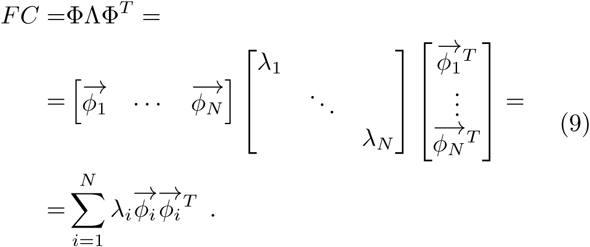

It follows that the magnitude of the eigenvalue gives a measure of the role of the corresponding eigenvector in reproducing the original matrix.

Taking into account all these observations, we identify the functional hub regions of the mouse brain as the regions with the largest eigenvector components, in absolute value, associated with the three largest eigenvalues of the FC matrix.

The code to run resting state simulation are code 1-2 and code 2-1 (Supplementary materials).

#### Modelling altered connectomes in pathological contexts

TVB allows manipulating the structural connectivity by selectively changing the strength of the connections between brain areas in order to mimic structural lesions. Using this tool, we have simulated mouse brain dynamics mimicking some aspects of the anatomical reorganization found in mesial temporal lobe epilepsy: the neuronal connection lost in hippocampal regions, in particular fields CA1 and CA3 (Esclapez et al., 1999). In order to reproduce this feature *in silico*, we have removed all the in-coming and out-coming connections of fields CA1 and CA3 of the hippocampus and then scaled all the connection strengths by a constant factor, so that the total weights of the modified SC is equal to the one of the original matrix. We simulated the resting state BOLD activity and we calculated the FCD matrix as described in the previous sections.

The code to run resting state simulation in pathological condition is code 3-1 (Supplementary materials).

### Epileptic spread in silico

#### The epileptic network node model

The Epileptor (Jirsa et al., 2014) is a model describing the onset (through a saddle-node bifurcation), the time course and the offset (through a homoclinic bifurcation) of seizures with 5 state variables that operate at 3 different time scales. The variable that guides the neural population through the bifurcations is the slow permittivity variable, *z*, which operates at the slowest time scale. Ensemble 1, comprising the variables *x*_1_ and *y*_1_, describes the fast discharges registered during ictal states and stable state observed during interictal states; it operates at the fastest timescale. Finally ensemble 2 (*x*_2_, *y*_2_) operates at the intermediate time scale and accounts for spike-and-wave events. The interaction between the variables of the system is the following: ensemble 1, trough the function *g*(*x*_1_), excites ensemble 2, which in turn inhibits ensemble 1 through *f*_1_(*x*_1_,*x*_2_); both the ensembles are coupled to the slow variable, and the first ensemble acts directly on *z*. Proix et al. (2014) propose a permittivity coupling between brain areas via a linear difference coupling function that links the fast subsystems with the slow variable *z* with the weights given by the distance *c_ij_*.

The full model equations read:

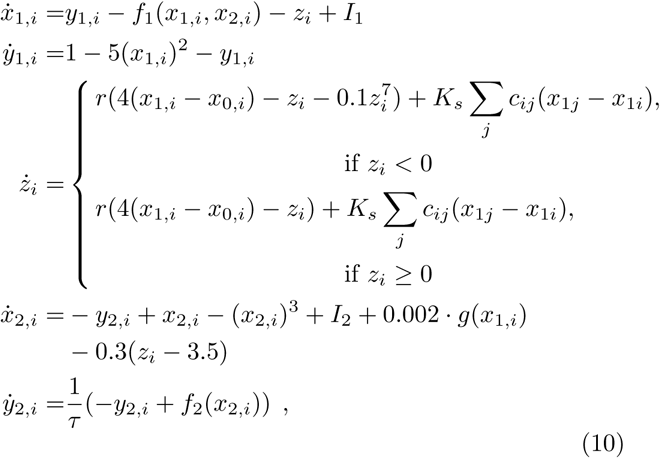

where

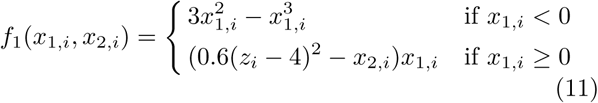

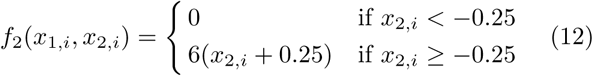

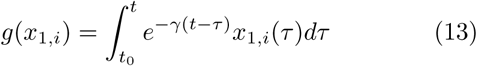
with *I*_1_ = 3.1, *I*_2_ = 0.45, *τ*= 10, *γ* = 0.01; the permittivity coupling term *K_s_* is fixed to -60.

The degree of epileptogenicity *x*_0_ of a brain region *i* is a parameter that establishes if the region generates seizures autonomously.

#### Integration scheme and epileptogenicity zone

The epileptogenic zones in the model are nodes of the network that are implemented in the simulation with an epileptogenicity value, *x*_0_, so that, for those nodes, the transition between the pre-ictal and the ictal state occurs spontaneously (Proix et al., 2014). An isolated brain area with *x*_0_ ≤ 2.06 is epileptogenic, otherwise the area is in its equilibrium state and it can generate seizures only if an external stimulus pushes it through the transition and makes it fall in the propagation zone (Proix et al., 2014). In the simulated mouse brain, the classification of brain areas in epileptogenic and propagation zone follows the experimental results of the work of Toyoda et al. (2013) in which the authors, using recording electrodes, evaluate the seizures propagation in rats with spontaneous seizures. The authors observe that the earliest seizure activity is recorded most frequently within the hippocampal formation and then spreads, in chronological order, in the subiculum, the entorhinal cortex, the olfactory cortex, the neocortex and the striatum; in 7 over 10 rats analyzed in the paper the epileptogenic region is likely identified in either hemisphere. Accordingly we set as epileptogenic, *x_0_* = −1.9 the left hippocampal regions (field CA1, field CA3 and dentate gyrus) and we set all other regions as propagation zones, *x_0_* = −2.1.

The differential equations of the model are integrated with the Heun stochastic method with an integration step equal to 0.04 ms; we use additive white Gaussian noise in the fast variables (*x*_2_ and *y*_2_) with mean zero and variance 0.0025. The signals are down-sampled to 1 ms. We set the pre-expression monitor in order to keep track of the local field potential, defined in the Epileptor as −*x*_1_ + *x*_2_, as well as the slow permittivity value *z*.

We define the time at which seizure initiates in a large brain region, as for example the olfactory cortex, as the mean of the seizure onset time of all the network nodes composing that region; in order to evaluate the chronological order of areas recruitment we define the seizure onset latency of a region as the difference between the time at which the seizure initiates in that region and the time at which the seizure has started in the epileptogenic zone, i.e. the hippocampal regions.

The code to simulate epileptic activity in mouse brain is code 4-1 (Supplementary materials).

## Results

### Virtualizing the mouse brain

#### Tracer-based connectome

In order to exploit present (and future) high-resolution structural information of the Allen Institute, we designed the Allen Connectivity Builder, a pipeline, which uploads their raw data and processes it in order to create a connectome and its brain volume representation. The user chooses four sets of parameters: the resolution of the tracing data (i); the way the connection strengths are calculated (ii); and the criteria used to include or not a given injected region based on its volume (iii) and its experimental significance (iv). The pipeline then computes automatically the averaged connection strength between any two regions. Since injections were only performed in the right hemisphere and since the mouse brain shows a high degree of lateral symmetry (Calabrese et al., 2015), the pipeline uses the mirror image to build the left hemisphere. If time delays are considered as an important variable to simulate whole brain activity, the length of each axonal tract becomes a key parameter. The Allen Connectivity Builder approximates the length of the tracts as the Euclidean distance between the centers of two regions. Finally, the pipeline automatically builds the brain volume using the same parcellation as used to build the connectome. TVMB includes a region volume mapping visualizer to display the brain volume as sections and the results of the computations in the brain sections.

An example of a structural connectivity matrix obtained through the Allen connectivity builder is shown in figure 1a, and the corresponding volume sections in figure 1d.

#### Diffusion MRI-based connectome

TVMB can also make use of user-based diffusion MRI data, enabling the virtualization of individual mouse brains. In order to use the analysis tools and the visualizer above, the brain volume should be uploaded in nifti format with the same parcellation as the connectome.

As an example, we have used here the high-resolution open-source mouse connectome of Calabrese et al. (2015) (figure 2a) which we have embedded in the Allen volume (figure 2d). In the general case, the user needs to upload the following files: (1) a weight matrix, i.e. a square matrix whose rows and columns label the areas in the parcellation and whose entry (*i, j*) represents the values of the connection strength between region *i* and region *j*; (2) a file containing the labels of the brain regions; and (3) the list of Cartesian triplets that specify the spatial location of each region (Sanz-Leon et al., 2015). As exhaustively explained in the TVB documentation (http://docs.thevirtualbrain.org/index.html), it is possible to provide additional information as the lengths of the tracts connecting the brain areas, or a file containing a vector providing a way of distinguishing cortical from subcortical regions, or the volumes where the connectome is embedded in nifti format etc.

**Figure 2:**
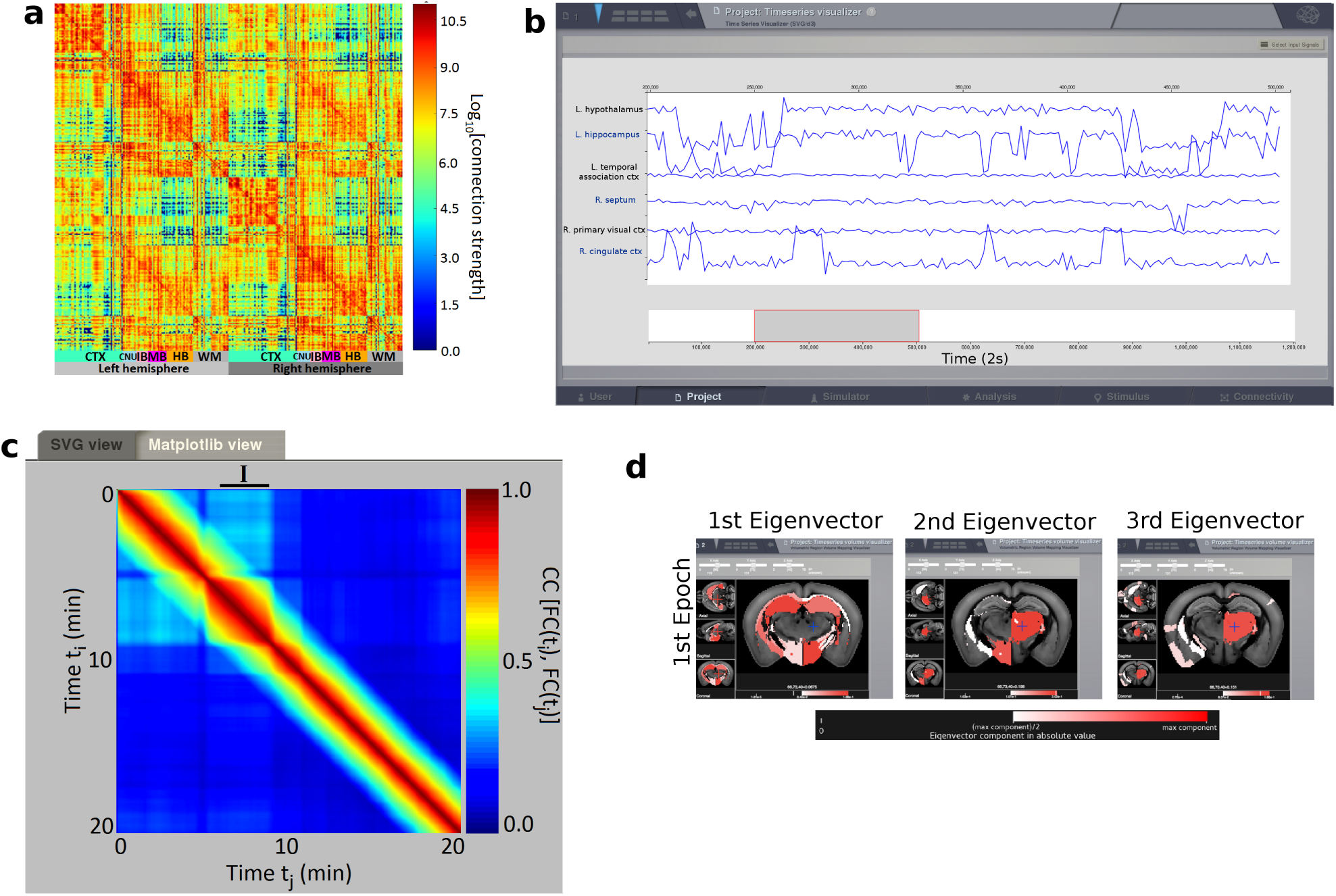
(a) Connectivity matrix obtained from Calabrese et al. (2015). (b) Simulated resting state BOLD time-series using the connectome shown in (a), and the eMFM to model the dynamics of each brain area. (c) FCD matrix obtained from the time-series. The black segment identifies the epoch of stability of the FCD identified with the spectral embedding technique. (d) Functional hubs detected in silico mapped in mouse brain sections using the brain region volume visualizer, as in figure 1c. The instructions and the codes to obtain the results in figure are respectively in Tutorial 1-1 and code 2-1 (Supplementary materials).

### Simulated brain activity

Once a virtual brain is constructed, the TVB environment generates a large-scale brain network equation (Jirsa, 2009) offering multiple ways to produce electrophysiological and neuroimaging signals, and analyze their dynamics. We present three illustrative examples on how mouse brain network simulations can be accomplished. The scripts and the data necessary to reproduce all the simulations and results presented here are in Supplementary materials.

#### Resting state activity in the “healthy” brain

Since recent studies highlighted the importance of studying Functional Connectivity Dynamics (FCD) (Allen et al., 2012, Hansen et al., 2015) and the functional hubs of rodents brain (Mechling et al., 2014, Liska et al., 2015) during resting state activity, we introduce an analyzer able to calculate the FCD and to extract the functional hubs (details of the algorithms in methods section).

We focus on the non-stationary nature of the fMRI functional connectivity (FC) in resting state observed both in humans (Allen et al., 2012, Chang and Glover, 2010)and in rodents (Keilholz et al., 2013, Liang et al., 2015). Thanks to the simulator tool of TVB we simulate the resting state activity using the reduced Wong Wang model (Wong and Wang, 2006) in the dynamical regime studied by Hansen et al. (2015). The model differs from previous resting state models (Deco and Jirsa, 2012, Deco et al., 2013) by having a richer dynamical repertoire for each brain region, which results in a greater number of attractors for the global system.

The BOLD signals and the corresponding FCD matrix are shown in figure 1b, and 1c respectively. The blocks around the diagonal of the FCD matrix correspond to time intervals during which the FC(*t*)s are strongly correlated; following the work of Hansen et al. (2015), we call these periods, epochs of stability. The FCD analyzer, using the spectral embedding algorithm, detects three epochs of stability (black lines in figure 1c) in the FCD matrix. As explained in the method section, it is possible to identify the central nodes of the *i*-th network (*i*=1,2,3), i.e. the functional hub regions of the *i*-th epoch, as the nodes linked to the largest components associated with the largest eigenvalues of the FC matrix computed over the *i*-th epoch. The functional hubs identified using this argument by the FCD analyzer are plotted in the mouse brain sections in figure 1d. It is possible to notice several analogies between the simulated functional hubs and the ones previously reported in literature as the hypothalamus, the visual and somatosensory cortex (Mechling et al., 2014) and the agranular insulare area, the cingulate and temporal cortex (Liska et al., 2015).

Brain activity can be simulated also in individual virtual mouse brain built from fMRI diffusion data as explained before. As an example, we uploaded the detailed diffusion MRI connectome from Calabrese et al. (2015) and simulated subsequent resting state activity of its BOLD signals (figure 2).

#### Resting state activity in epilepsy

TVMB can be used to assess the functional consequence of the anatomical reorganization that takes place in most, if not all, neurological disorders. Temporal Lobe Epilepsy is a prototypical example of neurological disorder with well-described anatomical alterations (Esclapez et al., 1999, Chen and Buckmaster, 2005) and functional reorganizations (Centeno and Carmichael, 2014).

Using the tracer-based connectome described above, we removed the connections from the hippocampal CA3 and CA1 regions known to be lost in some forms of medial temporal lobe epilepsy. The simulated BOLD and the corresponding FCD are in shown in figure 3.

**Figure 3:**
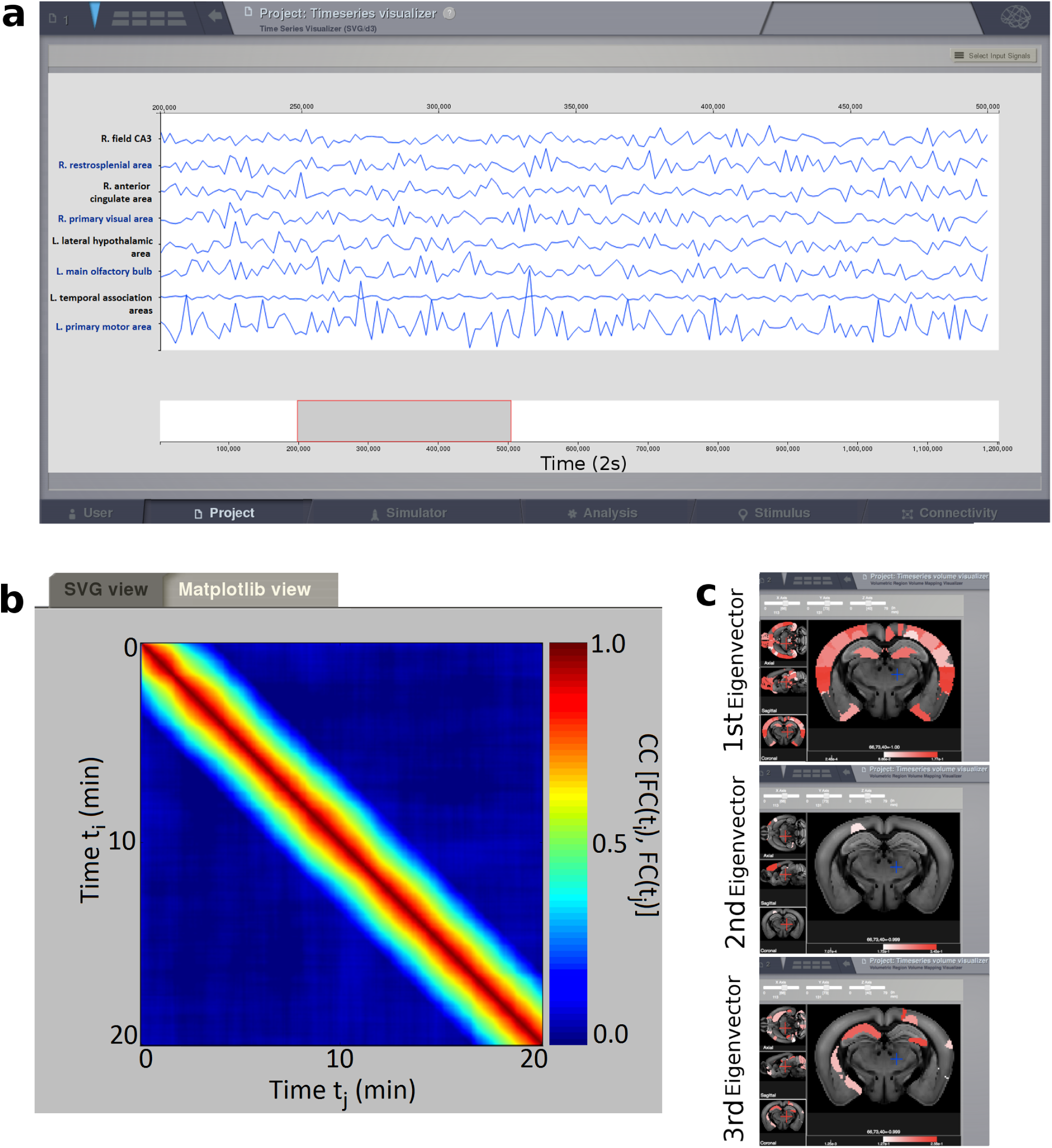
Figures (a) and (b) represent respectively the BOLD signals and the corresponding FCD matrix obtained by simulating the mouse brain in which some links are removed to mimic epilepsy conditions. (c) Functional hubs detected in the epileptic mouse brain after removing links as seen in some forms of epilepsy. The hubs displayed here are extracted from the FC matrix calculated over all the simulated BOLD signals (20 minutes), i.e. the global FC, since the FCD simulated in the epileptic mouse brain does not present evident sign of non-stationarity and consequently the epoch of stability can not be detected. The instructions and the codes to obtain the results in figure are respectively in Tutorial 1-1 and code 3-1 (Supplementary materials).

The comparison between the activity of the “healthy” and “epileptic” brain at the level of a single region, figure 1b and 3a respectively, does not provide any particular insight. However the differences in brain activity between the two conditions are revealed at the network level when computing the FCD (figure 3b). The functional connections that emerge in the “epileptic” brain are not correlated in time resulting in a suppression of the switching behavior of the FCD, as compared to the control connectome (figure 1c). As a result the functional hubs are modified. Since there is no switching, only hubs of global FC can be identified (figure 3c).

#### Seizure propagation

TVB also contains numerous models to generate EEG-like activity, including the Epileptor to simulate seizure genesis and propagation (Jirsa et al., 2014, Proix et al., 2014). As an experimental reference, we used the electrophysiological recordings performed by Toyoda et al. (2013) in a rat model of temporal lobe epilepsy. Based on the latter results, we used the left hippocampal regions as epileptogenic zones, and analyzed how and where seizures propagated in silico.

The results of the simulation are shown in figure 4. Each region is characterized by a specific time of seizure onset. The chronological order of the different areas recruited during seizure propagation is shown in figure 4b and 4c. The brain areas in abscissa in figure 4b are sorted according to the seizure onset latency rank found by Toyoda et al. (2013) in rats. Despite the difference in species (rat versus mouse), there is a remarkable analogy with experimental results, suggesting that the structural connectome (and the time delays it imposes) plays a key role in the spatiotemporal pattern of seizure propagation as already reported in humans (Jirsa et al., 2016, Proix et al., 2017).

**Figure 4:**
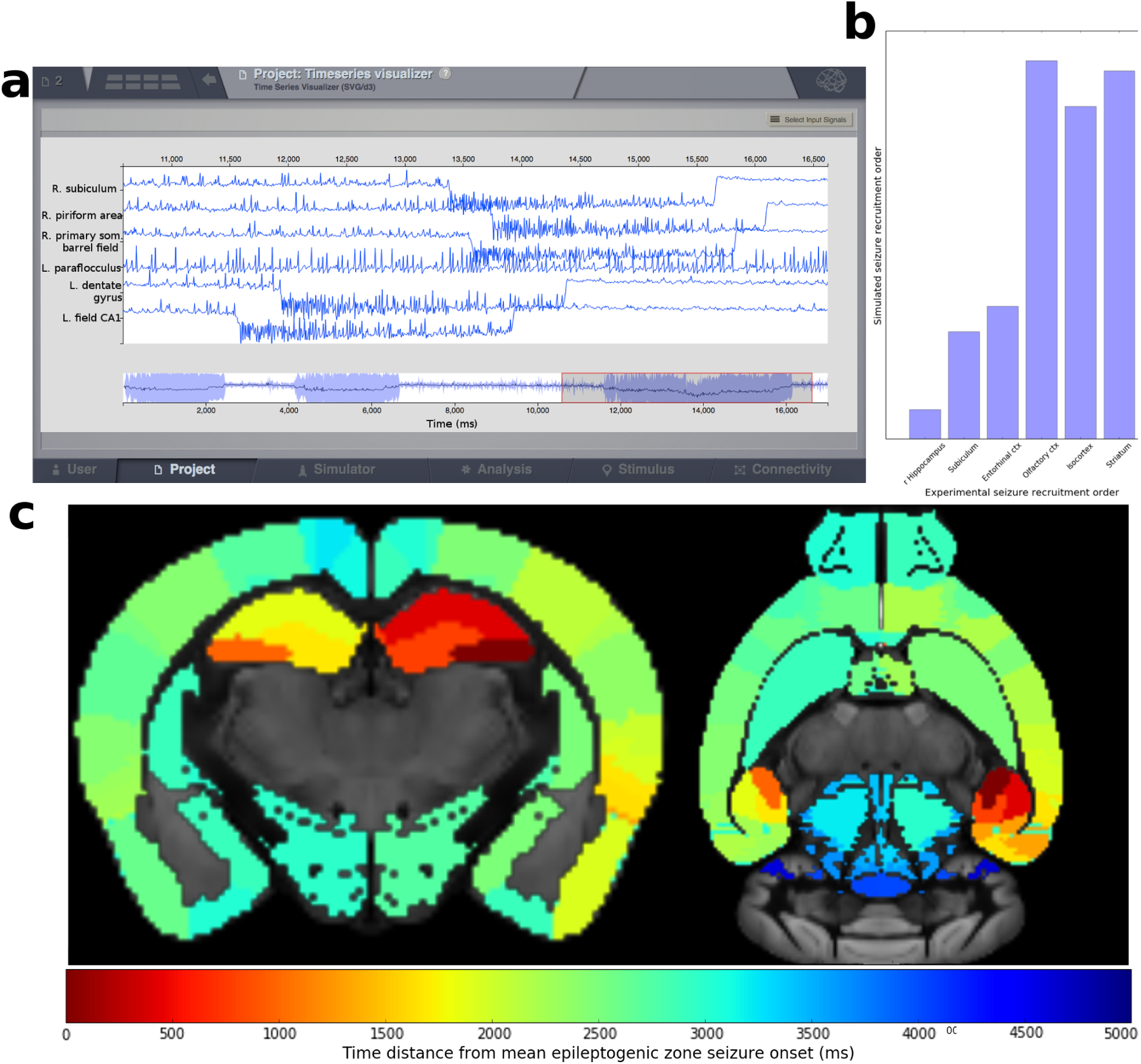
Simulating epileptiform activity in the mouse brain. (a) The time series show simulated seizure genesis and propagation (direct current recording) in silico. (b) The graph shows the propagation pattern. Time 0 corresponds to seizure onset in the left hippocampus. On the x axis, regions are ordered as they are progressively recruited in Toyoda et al. (2013). The y axis shows the average time of recruitment in arbitrary units of these regions after triggering a seizure in the left hippocampus in silico. Note the good match between simulated and experimental data. Extensive names of the region composing each group are illustrated in table 4-1 (Supplementary materials). (c) The time distance from seizure onset in the left hippocampus is given by the colorscale and plotted in the brain volume for each region. The instructions and the codes to obtain the results in figure are respectively in Tutorial 1-1 and code 4-1 (Supplementary materials).

### Interpreting and planning experiments with TVMB

#### Interpreting experimental data with TVMB

Physiological (e.g. normal aging) and pathological processes (e.g. neurological disorders) are associated with both structural (connectome) and functional (resting state networks) alterations. A central issue in neuroscience research is to understand how much structural alterations can account for functional ones. At present, both observations remain at the correlation level. In the case of aging, DTI and rsfMRI can be obtained at different times in a given animal (Figure 5). A virtual brain can be constructed at each time step, to simulate whole brain dynamics. Following data fitting, alterations found specifically at time *t* + 1 experimentally can be introduced in the connectome measured at time *t*. If the resulting in silico rsfMRI reproduces that experimentally measured at time *t* + 1, it is possible to conclude that these structural alterations are sufficient to explain the changes in whole brain dynamics.

**Figure 5:**
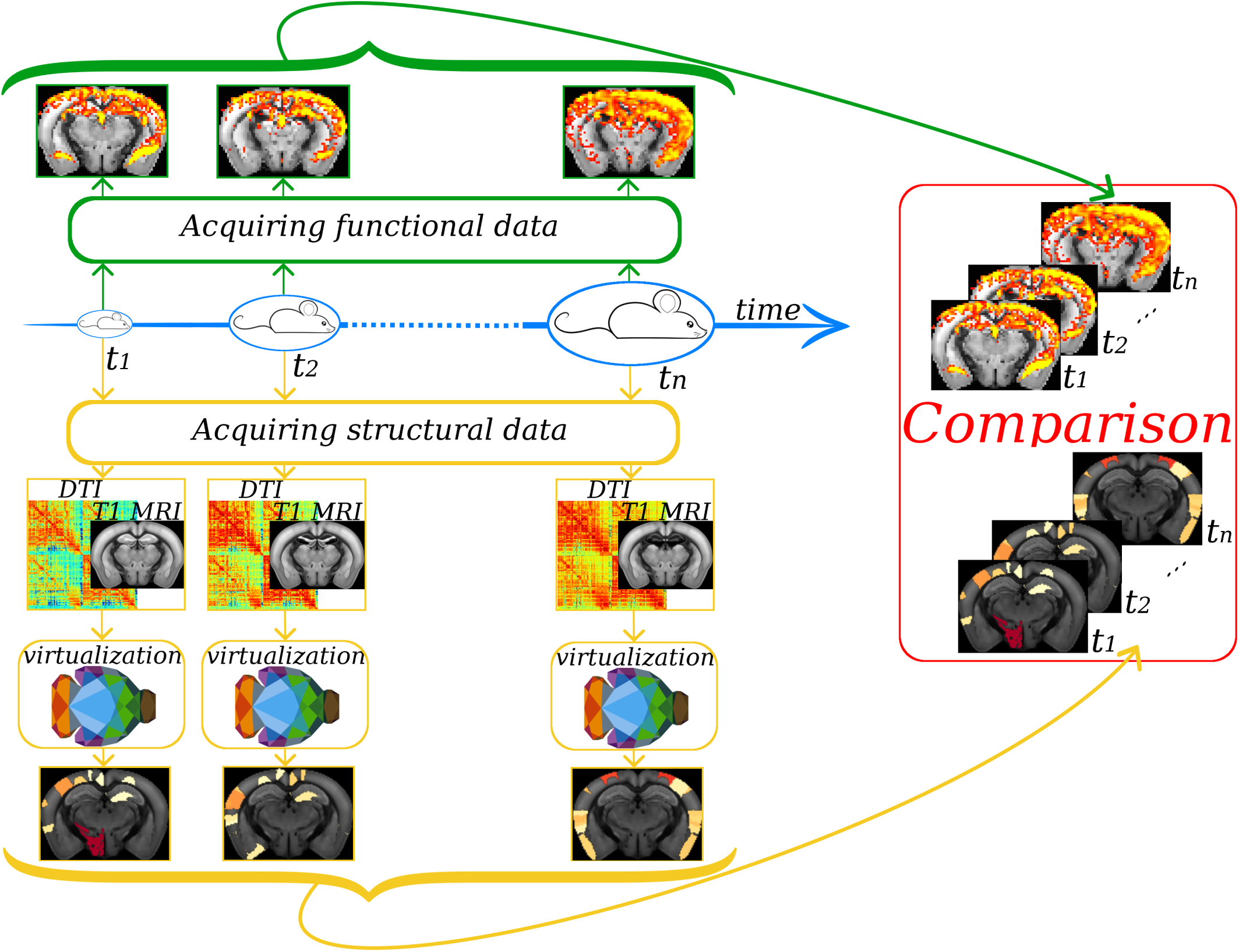
The cartoon illustrates how it is possible to use TVMB to do predictions when studying aging. A mouse can be scanned at different times t extracting anatomical and functional brain information. The anatomical information can be processed in order to obtain a connectome that can be used in TVMB to create a virtual mouse at each time step. The functional experimental information can be compared with the predictions done in TVMB, investigating how, for example, anatomical modifications during aging affect whole brain dynamics. Multiple other testable predictions can be done. For example, explore in silico which types of neurones can be stimulated (or silenced) to activate specific resting state networks. The predictions can then be tested in ad hoc transgenic mice with optogenetics.

#### Planning experiments with TVMB

We present two of the many possibilities offered by the platform. Brain surgery and stimulation are two common procedures used to treat patients, e.g. for epilepsy and Parkinson’s disease. After virtualizing a mouse model of these pathologies at a specific stage of their evolution, researchers can perform neurosurgery in silico and predict the efficacy of the procedure. Likewise, in silico stimulation of brain regions is straightforward in TVMB, which allows studying how resting state dynamics can be manipulated (Spiegler et al., 2016). The predictions thus generated can then be tested experimentally in vivo in the same mouse that was used to make them. Novel preclinical strategies may thus be tested in mice, before their possible clinical transfer.

Many brain functions require dynamical interactions and information transfer between numerous brain regions. The contribution of a given region is thus difficult to evaluate a priori. Using a parameteric study in TVMB, it is possible to predict which regions play a key role by successively activating and inactivating them. Then, one can plan the experiment, choosing the appropriate transgenic mouse line in order to control the identified region with optogenetics or pharmacogenetics. Such a priori knowledge provided by the in silico approach would considerably accelerate research.

## Discussion

TVMB opens a new set of research possibilities: it allows researchers, from different fields, to easily build specific/individual mouse brains (using various resolutions, weighting definitions and parcellations), to simulate different dynamical behaviors (using diverse neural population models, numerical integration schemes, and simulated neuroimaging modalities) and finally to analyze the results.

However, while TVMB is a highly generic framework, its underlying mathematical framework and simulation techniques make standard assumptions, among which the two most essential are that (1) the average activity of large populations of neurons is a meaningful quantification of the phenomena to be modeled and (2) the statistics of white matter fibers sufficiently describes how regions interact. Both resting state dynamics and seizure propagation, as demonstrated above, satisfy these assumptions. On the other hand, for example, fine grained spike timing effects would not well described within TVMB’s mathematical framework.

The wide range of possibility offered by rodent experiments will easily accommodate the validation of the parameterization required by all the modeling approaches contained in the software. This validation sometimes can proceed at a qualitative level for phenomenological models, such as the Kuramoto model of synchronization, but many detailed biophysical models allow for quantitive comparison with empirical data, such as spike timing (Brette and Gerstner, 2005). At the whole brain level, TVMB allows for direct comparison with common modalities such as EEG, MEG and fMRI, or common statistics thereupon such as functional connectivity; these comparisons allow for the characterization of parameter values in terms of their fit with empirical data and thus biological validity. The experimentally observed functional characteristics of the mouse brain (e.g. a functional hub during resting state or the effects of specific connections removal) can be easily imposed in the output of the virtual system, and through data fitting algorithms, it will be possible to retrieve the parameters of the model that give rise to that particular functional behavior. In this way, closing the circle, the reliability of the new predictions accomplished with the fitted parameter set will be improved; additionally the knowledge of the key features responsible of the different functional behavior allows to control and manipulate the system in silico, and, going a step further, also in vivo.

TVMB thus offers not only a conceptual framework to interpret neuroimaging data but, combined with experimental approaches, it also offers an operative framework to investigate the causal links between structure and function in the brain.

TVMB is an actively developed software, with new versions released regularly with new features. Among those targeted specifically for the mouse, the module which builds connectivities from the public Allen data will continue to evolve as the available dataset becomes richer. For example, when cortical layer annotations become available, it will be possible to construct mouse connectivities in which the cortical layers are distinct, allowing for example manipulations of inter-layer interactions. As dMRI protocols and tractography techniques become more established for rodent datasets, TVMB has potential to include modules which automate, with visual inspection across each step, the generation of connectomes for individual rodent data, directly from the DICOM slices provided by the acquisition equipment.

## Acknowledgement

We would like to express our gratitude to Calabrese E., Badea A., Cofer G., Qi Y., Johnson G. A., and the Allen Institute, for sharing their data with the community.

